# Effects of urbanisation, habitat characteristics, and management on garden pond biodiversity: findings from a large-scale citizen science survey

**DOI:** 10.1101/2023.12.08.570789

**Authors:** Zsuzsanna Márton, Barbara Barta, Csaba F. Vad, Beáta Szabó, Andrew J. Hamer, Vivien Kardos, Csilla Laskai, Ádám Fierpasz, Zsófia Horváth

**Author notes:** Corresponding author: Zsuzsanna Márton, HUN-REN Institute of Aquatic Ecology, Centre for Ecological Research, Budapest, Karolina út 29, 1113 Budapest, Hungary.

## Abstract

The rapid expansion of urban areas often leads to degradation, fragmentation, and loss of natural habitats, threatening biodiversity. While urban ponds might contribute substantially to the biodiversity of urban blue-green infrastructure, the role of garden ponds is still largely unkown. We lack a comprehensive understanding of how local habitat features, different forms of management, and urbanisation might impact the biodiversity of these habitats. This study aimed to reveal the importance of garden ponds via a country-wide online citizen science survey in Hungary, Central Europe. Data from over 800 pond owners revealed the occurrence and local frequency of various native animal taxa (amphibians, odonates, and birds), and introduced animals (e.g., fish). We collected data about pond features and management practices. We tested the effect of pond features, pond management, and landscape-level drivers (urbanisation, surrounding wetland coverage) on the presence of conspicuous animal taxa (adult amphibians and tadpoles, birds, odonates) to identify the potential drivers of the biodiversity of garden ponds. Key pond features including pond age, area, aquatic and shoreline vegetation were the most important factors, while algaecide addition was the most influential management practice negatively affecting amphibian presence. Urbanisation negatively affected the presence of adult amphibians and their tadpoles, but it was not associated negatively with the presence of odonates and birds. Our results indicate the high potential to utilise garden ponds as urban habitats surveyed with the help of the public. Developing effective urban biodiversity monitoring and conservation strategies are necessary for a better functioning blue-green infrastructure. The high level of engagement of pond owners, as in our survey, can create valuable data for achieving these aims.

## 1. Introduction

Land use changes including the intensification of agricultural activities, industrial development, and urbanisation are major drivers of species loss (Storch et al., 2022; Yang et al., 2024). Among them, urbanisation is a complex driver of global biodiversity loss (Concepción, 2021; Danneck et al., 2023; McDonald et al., 2019), acting through habitat loss, fragmentation, and degradation (Grooten & Almond, 2018; Stamenković et al., 2019; Tickner et al., 2020). It can create strongly selective environments e.g. via pollution (chemical light, noise), physical barriers obstructing organisms’ movement (roads and buildings), often leading to a limited species pool in urban habitats (Capotorti et al., 2023; Hill et al., 2021; Schueller et al., 2023). At the same time, urban habitats can also act as refuges for biodiversity depending on habitat planning and management (Soanes et al., 2019, 2023). Therefore, it is increasingly important to understand the role of local habitat features and management measures of the blue-green infrastructure in our efforts to efficiently conserve and increase urban biodiversity.

Ponds are increasingly recognised as key aquatic habitats, often supporting high levels of biodiversity within a region or landscape and hosting threatened or endangered species (Céréghino et al., 2007; Williams et al., 2004). Lentic aquatic habitats less than 5 m deep and ranging 1 m^2^ and 2-5 ha in size are generally considered ponds (Biggs et al., 2005; Céréghino et al., 2007; Richardson et al., 2022). It has been estimated that freshwater ponds (< 1 km^2^) cover more than 3% of the Earth’s surface, and constitute the most widespread type of inland standing waters (Downing et al., 2006). They often represent the most abundant aquatic habitats locally in natural landscapes and even in areas with high agricultural activity (Oertli et al., 2009; Ruggiero et al., 2008).

Garden ponds are popular features that can provide a range of ecosystem services including microclimate regulation, supporting biodiversity, and cultural services such as wildlife observation or connecting with nature (Hill et al., 2021; Oertli et al., 2023; Peeters et al., 2023). Yet, very limited information exists about the exact number and ecological roles of garden ponds in cities (Hill et al., 2021). Garden ponds can host threatened taxa like amphibians (Hassall, 2014), which are among the most vulnerable freshwater organisms worldwide (Hamer & McDonnell, 2008). At the same time, garden ponds may also act as a source of invasive species due to commercial trade, limited knowledge, and aesthetic preference thereby threatening native biodiversity (Patoka et al., 2017). To promote efficient management practices that support native species while minimizing the facilitation of invasion events, we need better understanding how current management practices affect the wildlife of these urban habitats. Currently, this represents a general knowledge gap likely linked to the novelty of this ecosystem type in urban blue-green infrastructure and also their limited accessibility to researchers.

Citizen (or community) science, i.e., the involvement of members of the public in scientific research and data production, is becoming a valued and necessary research tool for scientists to evaluate or gain knowledge about a wide range of scientific topics (Fraisl et al., 2022). Data collected by citizens and later harmonized and validated by experts can greatly improve our understanding of species diversity at large spatial and temporal scales as well as ecological dynamics and the conservation status of species (Steven et al., 2019). Garden ponds are difficult to access or sample as they are situated on private properties. With the involvement of their owners, citizen science data collection can not only be achievable, but is essentially the only feasible way to collect large-scale datasets from these ecosystems. Hence, for a better understanding of the biodiversity, ecology, and management of garden ponds, citizen science is an essential tool, which can at the same time lead to a high level of public engagement (Hamer et al., 2024; Peeters et al., 2023).

To improve our understanding on the ecological roles of garden ponds, we conducted a countrywide online survey in Central Europe (Hungary). The specific aims of our study were to 1) explore how frequently wild animals visit or inhabit garden ponds focusing on taxonomic groups that are conspicuous enough to be easily and reliably identified by the public (amphibians and their tadpoles, birds, odonates); 2) investigate the frequency of animal introductions, including potentially invasive ones, to garden ponds (i.e. animals such as fish, mussels, snails or crayfish introduced by the pond owner); 3) identify the potential drivers of native biota by testing the reported presence of animals against urbanisation. local pond features and management practices applied by the owners (for the latter, data was available from Hamer et al., 2024), with the expectation that urbanisation will have an overall negative effect on the presence of most taxa.

## 2. Materials and Methods

### 2.1. Citizen science survey

An online survey within the citizen science project “MyPond” was launched in 2021 to investigate the biodiversity, local features, and management practices of garden ponds in Hungary (Central Europe). The link to the questionnaire on the “MyPond” website (https://mypond.hu/en/) was distributed via traditional media outlets and social media. The survey consisted of a total of 30 questions: 6 multiple choices, 10 short answers, and 14 closed (yes-no) questions. We collected data on the local features of ponds, including age, area, substrate of the pond (concrete, rubber pond liner, plastic, and natural, the latter including soil, gravel, clay etc.), and the presence/absence of aquatic vegetation, shoreline vegetation, and fish. Local management data included circulation, cleaning, draining, leaf removal, and use of algaecide. The results from these management data and local pond features were summarised and published by Hamer et al., 2024.

The biodiversity dataset consisted of four animal taxa, amphibians (here, we refer to adult amphibians or post-metamorphic juvenile sightings) and their larvae referred to as tadpoles (reported separately from adult amphibians), birds, and odonates (we used the Hungarian term „szitakötő” which does not differentiate between damselfiles and dragonflies). The survey contained a link to the amphibian identification key (illustrated with drawings) of the BirdLife Hungary to help the citizens correctly identify amphibian species (MME (2022). https://www.mme.hu/en/node/4440), which is available as a free smartphone application. The identification key has been developed especially for citizen science monitoring and it guides a non-scientist volunteer through the most important features (including shape, colour, and habitat) of amphibians, resulting in species-level identification. Also, there was an option to name the species of birds they had seen in their gardens.

### 2.2. Data validation and processing

The questionnaire is still available on the website and collects data for future purposes. Between 2021 and 2022 (6 July 2021 - 9 September 2022), 834 participants filled out the questionnaire, of which a validated subset of 753 responses was used for statistical analysis (Figure 1). Validation was carried out to exclude duplicates and incomplete data. Duplicates were identified as having the same geolocation or email address provided by the owners. However, respondents may have multiple ponds in their garden, therefore the length, width and depth were also inspected before exclusion. If the pond sizes were roughly the same, it was considered a duplicate. We also excluded ponds with an area over 200 m^2^, which we set as a threshold based on the frequency distribution of pond areas (the largest was 24000 m^2^, but 98% ponds were smaller than 200 m^2^). We approximated the pond surface area based on the lengths and widths (that pond owners provided), by the multiplication of these measurements. Pond age was used as 0 if the pond was created in the year when the survey was filled out. In some cases, pond age was missing from the dataset, since the previous homeowner constructed the pond, and the current owner did not hold specific information on the exact year of construction. If they provided the information of when they moved in, we calculated the pond age as the number of years since then.

**Figure 1.**
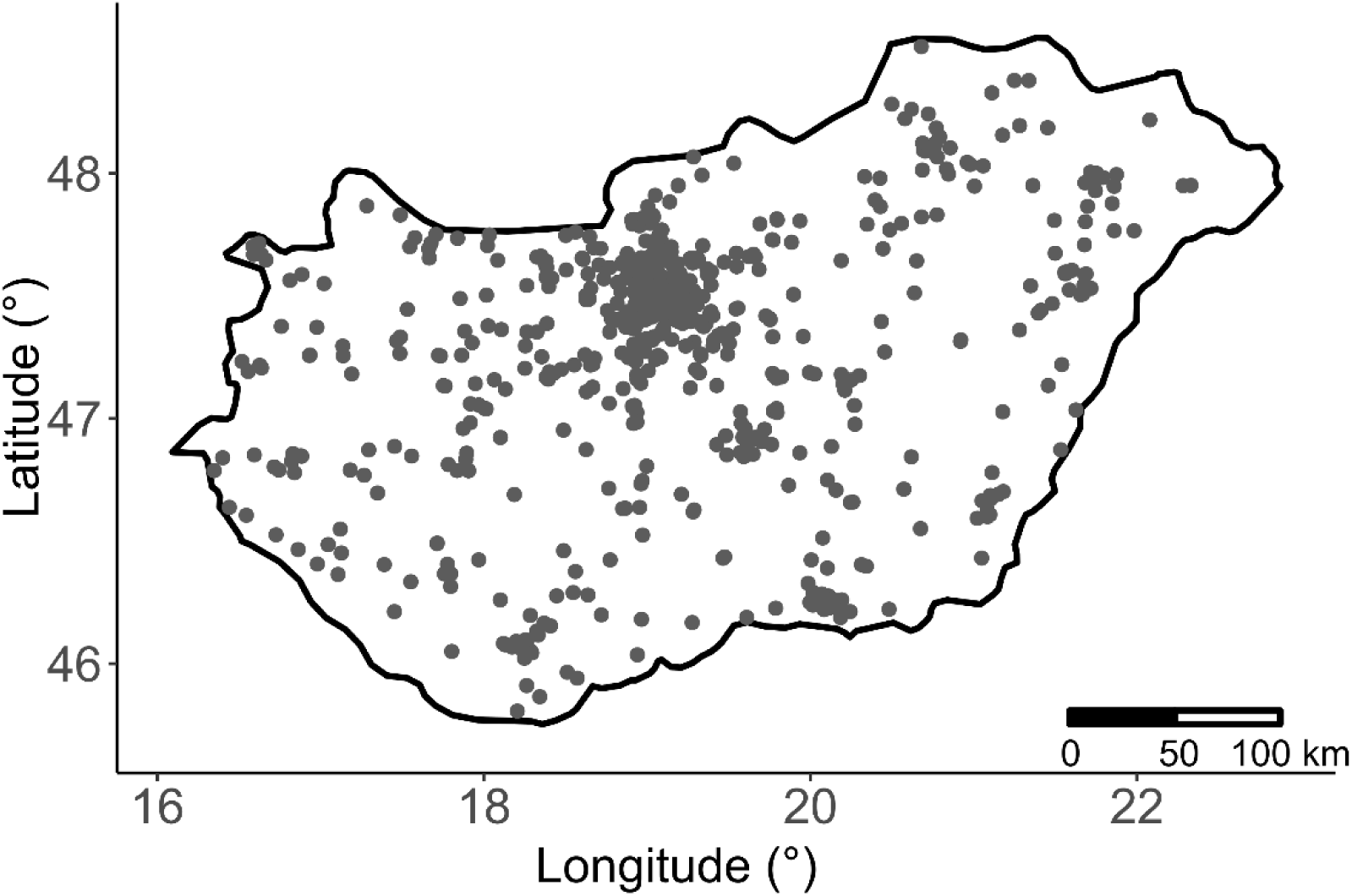
Location of the 753 garden ponds in Hungary used in the analyses. (Map was created with R v. 4.2.2)

In the case of the amphibian species dataset, we excluded amphibian species if pond owners made statements such as “we have not seen that species in a few years”. However, we included species if the owner reported them as “rarely” seen. Furthermore, entries were excluded from the amphibian species dataset if it only included the name “frog”, “toad”, “newt”, or “tadpoles”, as species identity could not be determined. “Green frogs” were grouped into the *Pelophylax* spp. complex as the three species that comprise this group are morphologically difficult to separate, resulting in an amphibian dataset of 332 entries. Rare species were also cross-checked with the most up-to-date species distribution maps (https://herpterkep.mme.hu/index.php?lang=en) and retained only if their occurrence had already been proven within the region of the sighting. Regarding birds, due to the high number of potential bird species, we decided to group the reported birds from the open-ended question (Table S1) into larger categories: songbirds, wading birds, swallows, woodpeckers, ducks, and pigeons. Here, songbirds included thrushes, sparrows, finches, starlings, tits, larks, and nuthatches while wading birds included herons, egrets and storks. These larger categories can be still reliably identified by the public and were actually used by many of the participants in their original responses to this question where they could not fully identify the species they had seen. For the analysis on the potential drivers of occurrence, we further grouped these data into three main categories according to their pond use type (see below). Data were excluded if respondents did not specify the species or categories, resulting in a dataset of 661 entries. The full questionnaire can be found in the supplementary information (Table S1).

The resulting datasets included both occurrence (binary; 0, 1) and local frequency (ordinal scale; 0, 1, 2, 3) data of each recorded animal taxa (amphibians and their tadpoles, odonates, and birds). Pond features (pond age, area, substrate, aquatic vegetation, shoreline vegetation, and fish) and management (cleaning, draining, leaf removal, and the use of algaecides) were used as binomial and ratio scale data in the statistical analysis (Table 1).

**Table 1.** Data types in the validated dataset.

| **Main variable groups** | **Recorded variables** | **Data type** |
| --- | --- | --- |
| **Biodiversity - occurrence** | **amphibians and their tadpoles, birds, odonates** | Binomial (0, 1) |
| **Biodiversity - local frequency** | **amphibians and their tadpoles, birds, odonates** | Ordinal (0, 1, 2, 3, corresponding to no, singular, occasional, and frequent sightings per pond) |
| **Pond management** | **circulation, cleaning, draining, leaf removal, algaecide** | Binomial (0, 1) |
| **Pond features** | **age, area, substrate (concrete, rubber, plastic, natural), aquatic vegetation, shoreline vegetation, fish presence** | Ratio scale and binomial (0,1) data |

### 2.3. Landscape-level drivers: urbanisation and wetland coverage data

The area of surrounding urban land and wetland coverage, predicted to be landscape-level drivers of species occurrence, were calculated using the Ecosystem Map of Hungary (project KEHOP-430-VEKOP-15-2016-00001, Ministry of Agriculture, 2019). The wetland coverage category contained pixels of wetlands, marshlands, temporary wetlands, standing waters, and flowing waters, while the urban land category (hereafter urbanisation) contained pixels of tall buildings, short buildings, sealed roads, dirt roads, railways, other artificial surfaces, and urban green spaces with and without trees. The number of raster pixels (20×20 m) was counted within a 1 km radius around each focal pond using the Zonal Histogram tool (QGIS Development Team, 2022). A 1 km buffer was chosen as a widely used measure for studying birds (Humphrey et al., 2023) and it also contains the mean and maximum dispersal distances of most amphibian species expected to occur in Hungary (Kovar et al., 2009). While we acknowledge that this might have created some overlap in the individual buffer zones of some of the ponds (mostly in Budapest where ponds were situated closer to each other), given the average pond distance of 3514 m in the dataset, it still provided a wide and largely independent gradient of landscape cover for most of the studied ponds. While a few ponds were less than 1 km apart, resulting in partially overlapping buffer zones, this was only the case in 0.09% of all the pond pairs, hence the effect of this should have been minimal. The classification of land coverage was used as defined by the Ecosystem Map of Hungary (Ministry of Agriculture, 2019). Urbanisation and wetland coverage were then calculated as the proportion of total pixels within the 1 km radius covered by the respective land cover type (Oertli & Parris, 2019). All spatial analysis was carried out in QGIS 3.28.1 (QGIS Development Team, 2022).

### 2.4. Statistical analysis

To visualise the occurrence and the local frequency of animal taxa, we created barplots and pie charts using the “ggplot2” R package (Wickham, 2016).

To examine the relationship between amphibian community composition and pond features, pond management, and landscape-level drivers (urbanisation and wetland coverage), we performed canonical correspondence analysis (CCA) using the “vegan” R package (Oksanen, 2017). Prior to the CCA analysis, we excluded ponds with no amphibian species recorded, and the analysis was based on the amphibian species occurrence data (binomial). To normalize the distribution of variables belonging to pond features and landscape-level drivers, we used log transformation for pond area, and square root transformation was applied for wetland coverage prior to the CCA. Pond management variables were included as binomial data. Significant explanatory variables among pond features, management, and landscape-level drivers were identified based on an automatic stepwise (backward and forward) model selection procedure using permutation tests (“ordistep” function, direction = “both”, n = 999 permutations).

To analyse which pond features, management types, or landscape-level drivers had the strongest effect on the occurrence of each four investigated animal taxa (amphibians, tadpoles, birds and odonates; Table 1), generalised linear models (GLM) were fitted with a binomial distribution and logit link function (hereinafter logistic regression model). Model selection was carried out based on Akaike’s Information Criteria (AIC) with the “MASS” R package (Venables & Ripley, 2002). In the case of birds, we also ran separate models where bird categories were additionally grouped into three main groups according to the way they use the ponds: wading birds (herons, egrets, and storks), ducks, and other birds (mostly including songbirds that mainly use the ponds for drinking). To test for potential spatial autocorrelation in the resulting final GLMs, we used the Moran’s I test from the „DHARMA” R package (Hartig, 2022). As no significant autocorrelation was found (Table S2), we kept the original GLM models for the analyses on the effects of pond features, management types, and landscape-level drivers. Visualisation was carried out with the “ggplot2” (Wickham, 2016) and “ggbreak” (Xu et al., 2021) R packages.

All statistical analyses and illustrations were made with the R program v 4.2.2 (R Core Team, 2022).

## 3. Results

Based on the occurrence data reported by 753 respondents, odonates (in 96.5% of the ponds) and birds (96.5%) were observed at almost all garden ponds. While the occurrence of amphibian sightings (81.4%) was higher than that of tadpoles, the latter were still observed in close to two-thirds of the ponds (62.1%). Among the amphibian species, *Pelophylax* spp. was the most frequently reported species (29.9%). The most frequently reported birds were songbirds (88.3%, including tits, blackbirds, sparrows, etc.). Within the introduced animal taxa, fish were the most frequent (84.4%), while snails, turtles, mussels, and crayfish were all introduced to less than 6% of the ponds (Figure 2).

**Figure 2.**
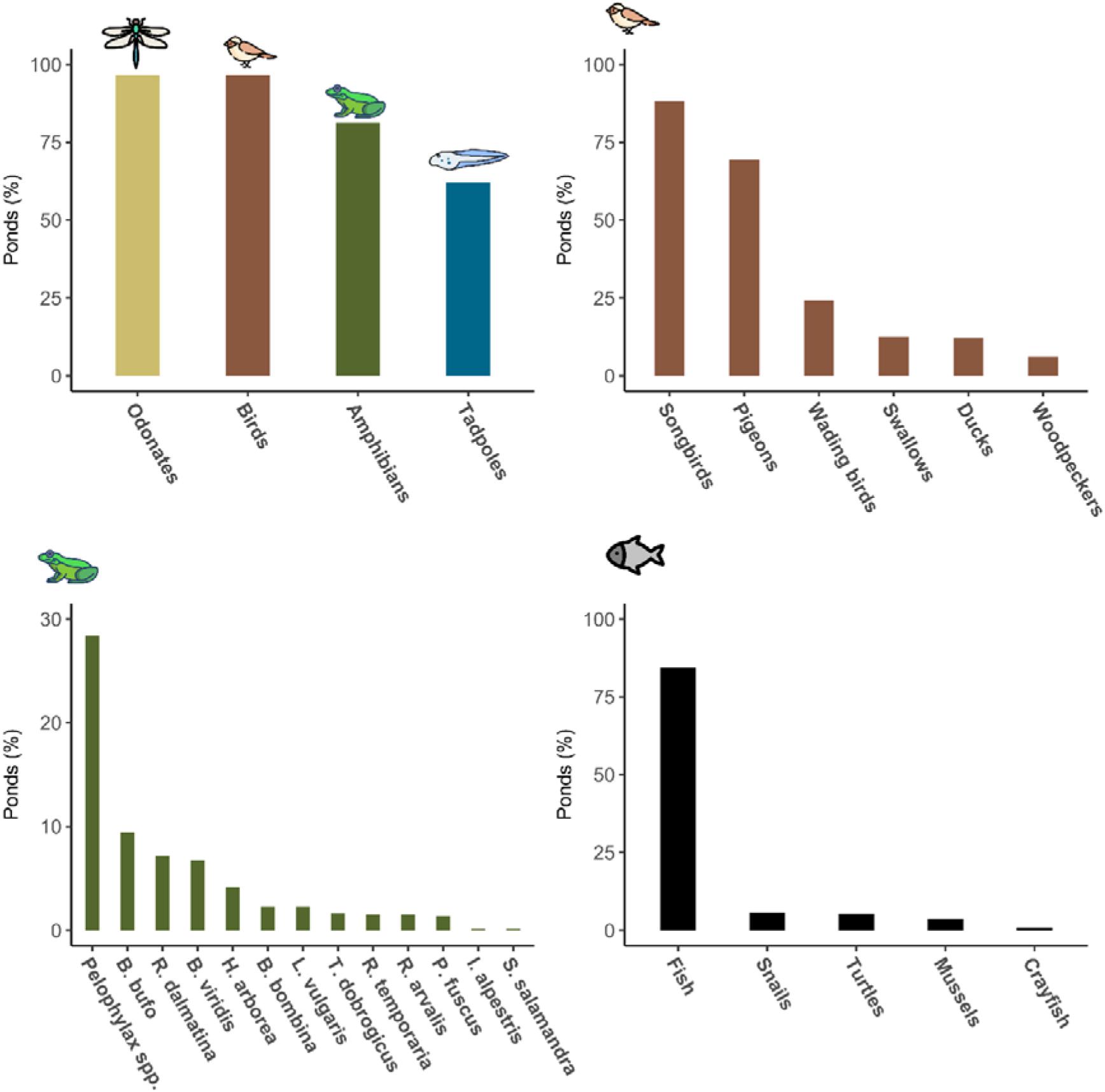
Percentage of all ponds with the occurrence of the animal taxa reported by the citizens. Samples sizes were n=667 in case of types of birds (top right), and n=753 in the other three datasets. For the abbreviation of amphibian species (bottom left figure), see the legend of Figure 4.

Local frequency data of amphibians, birds, and odonates revealed that most (60%, 78.5%, and 86.5%, respectively) of the respondents regularly see these animal taxa in their ponds. Among them, tadpoles were absent from the highest share of ponds (37.8%), but amphibian breeding was regularly observed in over 40% of the ponds detected via tadpole presence (Figure 3).

**Figure 3.**
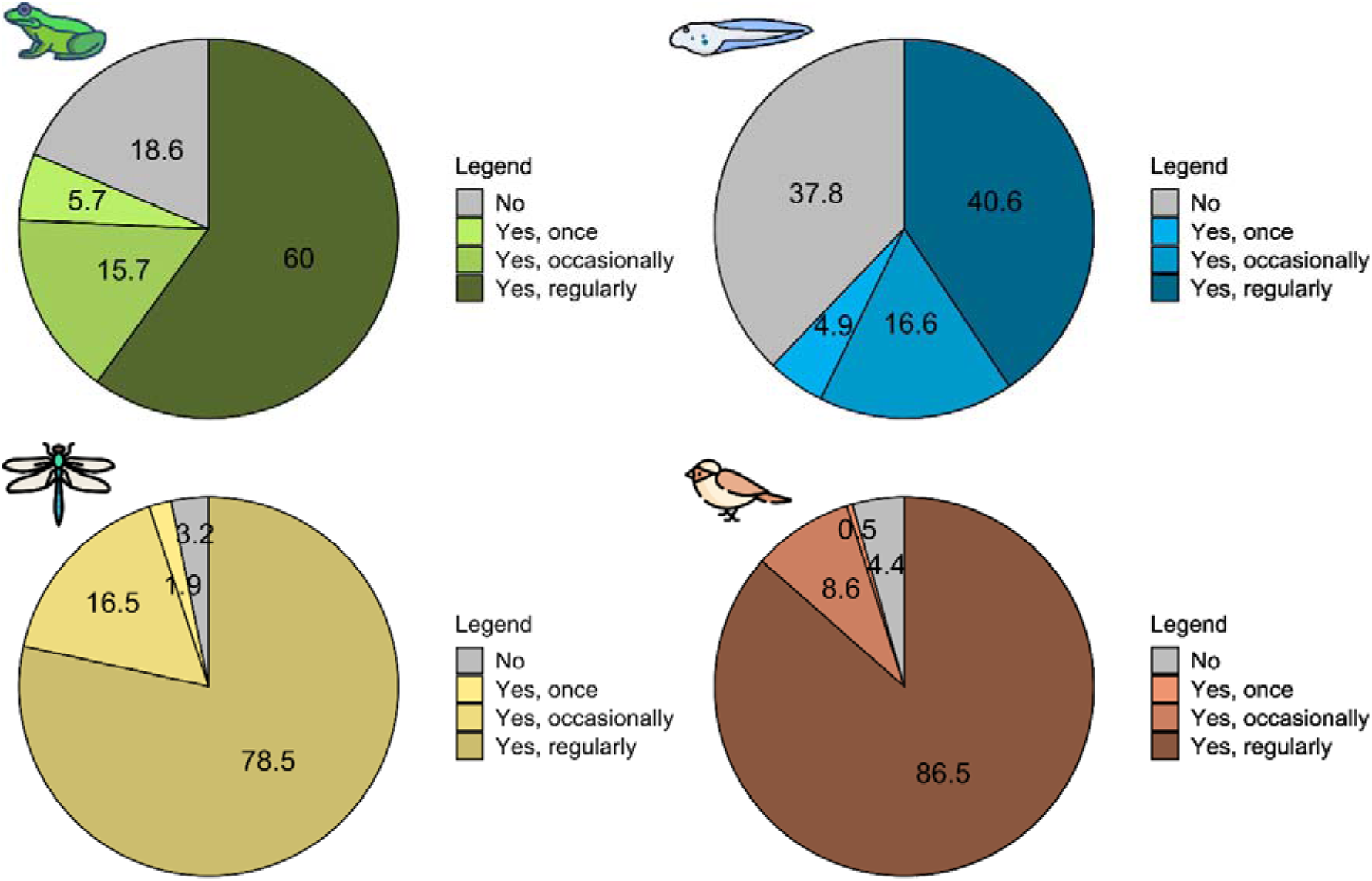
Share (%) of each response on the local frequency of animal taxa (amphibians and their tadpoles, odonates and birds) in garden ponds (n=753)

Participants recorded the presence of 13 amphibian species in garden ponds: three newt species (*Ichthyosaura alpestris*, *Lissotriton vulgaris*, *Triturus dobrogicus*), nine anurans (*Bombina bombina*, *Bufo bufo*, *Bufotes viridis*, *Hyla arborea*, *Pelobates fuscus*, *Pelophylax* spp., *Rana arvalis*, *R. dalmatina*, *R. temporaria*) and one salamander (*Salamandra salamandra*). Amphibian species composition (based on occurrence data, Table 1) was significantly influenced by pond age (p = 0.050), pond area (p = 0.043), rubber pond liner (p = 0.034), shoreline vegetation (p = 0.072), algaecide usage (p = 0.077), the introduction of fish (p = 0.001) (Figure 4). There was a negative association between *Rana arvalis* and pond age, i.e., it occurred mostly in newly installed ponds with rubber pond liner and shoreline vegetation. *Lissotriton vulgaris*, *Rana temporaria,* and *R. dalmatina* responded negatively to fish presence and algaecide treatment. *Bufotes viridis* was positively associated with urbanisation. *Salamandra salamandra*, *Ichthyosaura alpestris, Triturus dobrogicus, Bombina bombina, Hyla arborea,* and *Pelobates fuscus* were positively associated with large, well-vegetated ponds with a natural substrate, which are only occasionally drained (Figure 4).

**Figure 4.**
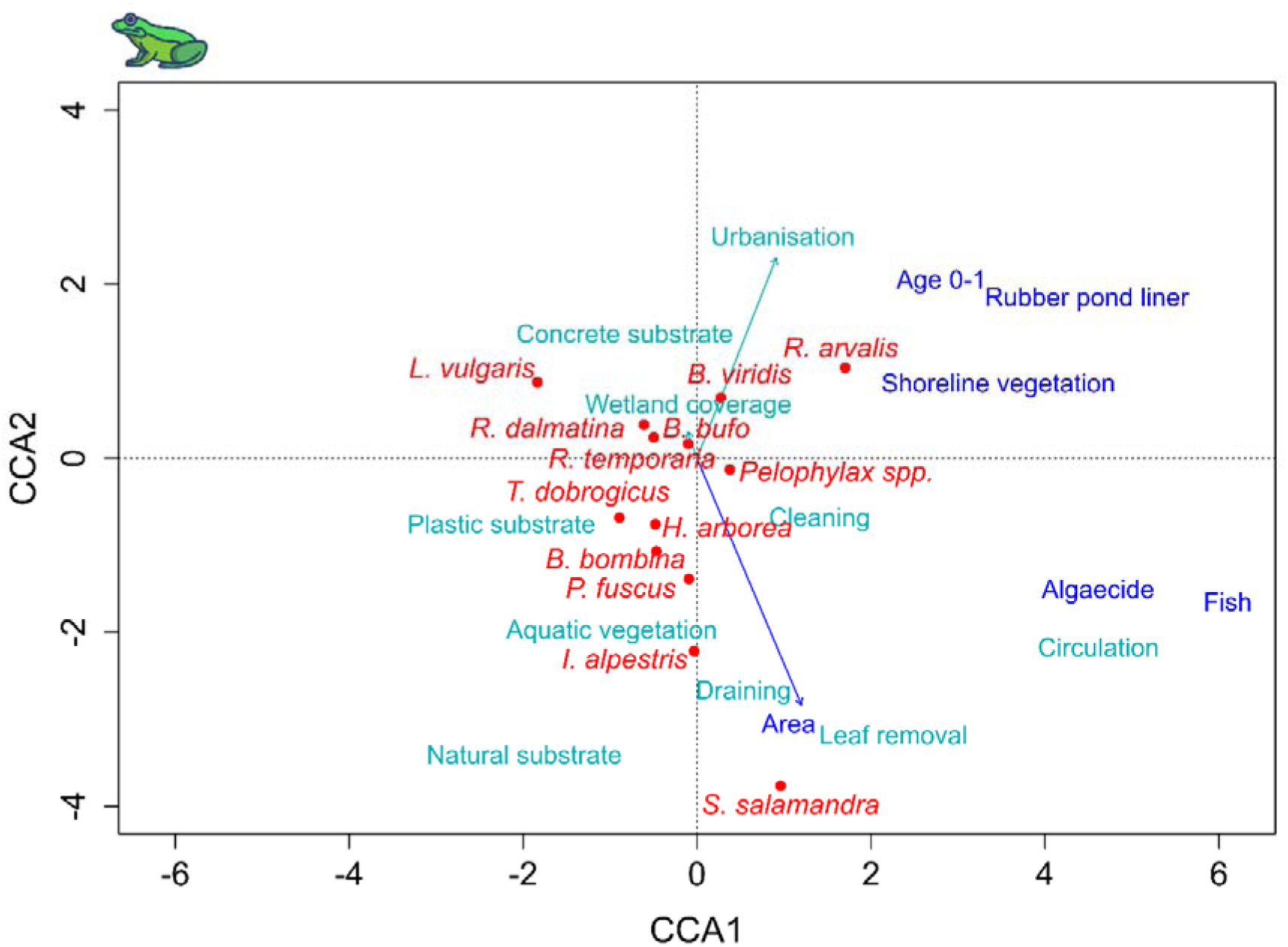
Canonical correspondence analysis (CCA) biplot based on the reported prevalence of amphibian species, environmental predictors, and level of urbanisation (red points: species scores, dark blue arrows: significant predictors, light green arrows: all other predictors). The analysis is based on data from 332 ponds with validated species data. (Abbreviated species names: *I. alpestris = Ichthyosaura alpestris*, *L. vulgaris* = *Lissotriton vulgaris*, *T. dobrogicus* = *Triturus dobrogicus*, *B. bombina* = *Bombina bombina*, *B. bufo* = *Bufo bufo*, *B. viridis* = *Bufotes viridis*, *H. arborea* = *Hyla arborea*, *P. fuscus = Pelobates fuscus*, *R. arvalis* = *Rana arvalis*, *R. dalmatina* = *Rana dalmatina*, *R. temporaria* = *Rana temporaria, S. salamandra* = *Salamandra salamandra*)

Based on the results of the logistic regression models, urbanisation was negatively associated with the presence of amphibians and their tadpoles, while it generally had a positive effect on the presence of odonates and birds (Figure 5, Table S3). Wetland coverage did not have a significant effect in any of the models, although a positive trend was found for amphibians (Figure 5, odds ratio (OR) = 34.300, p = 0.109). The most frequently significant pond feature was pond age (logistic regression model of amphibian: OR = 0.256, p < 0.001; tadpoles: OR = 0.351, p < 0.001; odonates: OR = 0.154, p < 0.001; birds: OR = 0.155, p < 0.001), followed by rubber pond liner (amphibians: OR = 2.150, p = 0.004; tadpoles: OR = 1.997, p = 0.003; odonates: OR = 2.510, p = 0.048; birds: OR = 2.510, p = 0.048; wading birds: OR = 2.655, p = 0.021) and pond area (amphibians: OR = 1.036, p = 0.001; tadpoles: OR = 1.014, p = 0.001). Amphibians and their tadpoles, odonates, and birds were less likely to be present in or at newly installed ponds (0-1 year). Pond area had a positive, while algaecides had a negative, effect on the presence of both amphibians and their tadpoles. There was a positive effect of aquatic vegetation on tadpoles, odonates and birds, however, shoreline vegetation was only significant for the presence of tadpoles (Figure 5, Table S3).

**Figure 5.**
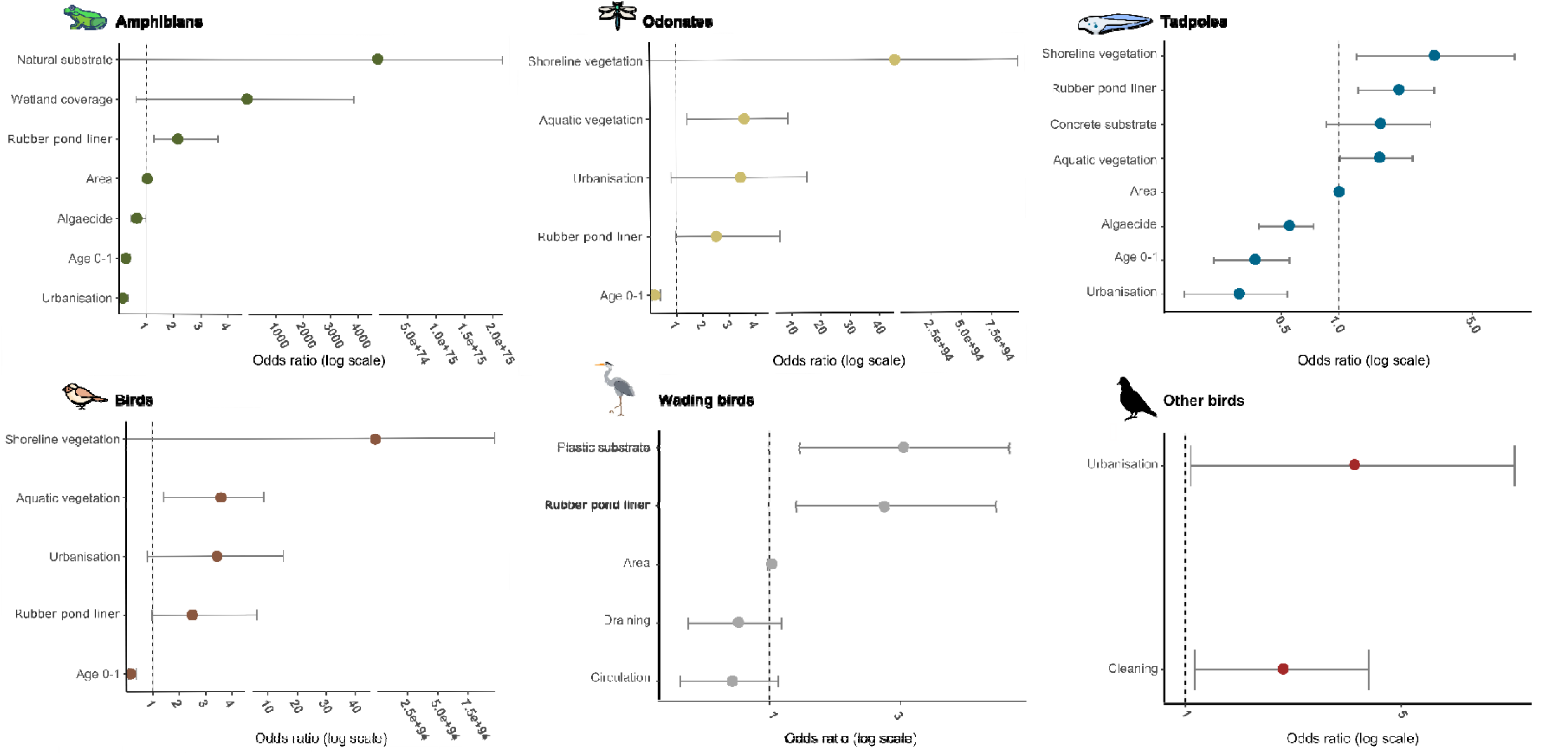
Summary of the best-fit logistic regression models (based on AIC) odds ratio (filled circles) and their 95% confidence intervals (whiskers) testing the relationship between the occurrence of animal taxa and pond features and management. Samples sizes were n=753 for amphibians, odonates, birds, and tadpoles, and n=667 for other birds and wading birds. (Birds: all birds; Wading birds: herons, egrets and storks; Other birds: songbirds, pigeons, woodpeckers

## 4. Discussion

Our citizen science survey provided comprehensive large-scale data on the presence of multiple animal taxa of garden ponds, covering an entire European country. Odonates and birds were observed at almost all 753 garden ponds, but amphibians and their tadpoles were also common. Local frequency data revealed that most respondents regularly see these animal taxa. Urbanisation had an overall negative effect on the presence of amphibians and tadpoles; conversely, odonates and birds showed an unexpected positive association with it. The age of the ponds was a general predictor, with animals less likely to be present in newly installed ponds. Among local pond features, pond area and rubber pond liner were the most important across the animal taxa, mostly with positive effects. Among management types, algaecide addition had a negative effect on amphibians and their tadpoles. Our data also highlighted that owners introduce various animal taxa into their ponds, most frequently fish, with potential consequences for the native biodiversity surrounding garden ponds. Our results overall illustrated the potentially important role of garden ponds in urban biodiversity and conservation by providing secondary habitats for multiple animal taxa.

### 4.1. Effect of urbanisation

Urbanisation had disparate effects across the animal taxa we investigated. It had a negative effect on the occurrence (presence/absence) of amphibians and their tadpoles, in line with our original hypothesis and also corroborating earlier findings (Hamer & Parris, 2011; Rubbo & Kiesecker, 2005). Amphibians are particularly vulnerable to urbanisation and fragmented landscapes, comprised of roads, houses and buildings that can impede dispersal between ponds and natural upland habitats that are important for amphibians to complete their complex life cycle (Hamer & McDonnell, 2008; Scheffers & Paszkowski, 2012). Although urbanisation generally has a negative effect on amphibian diversity, some species are urban adapters (McKinney, 2002), favored by larger body sizes and the ability to persist even in fragmented landscapes (Hahs et al., 2023). For example, *Bufotes viridis* was positively associated with urbanisation in our study. This species has a relatively large body size and persists in the centre of European cities and towns (Landler et al., 2023; Mazgajska & Mazgajski, 2020; Sistani et al., 2021). The *Pelophylax* spp. complex had no discernible associations with any of the variables, probably because these species are habitat generalists and can maintain metapopulations in both natural and urban landscapes (Herczeg et al., 2012; Ficetola and De Bernardi, 2004). Conversely, species such as *Ichthyosaura alpestris* which is associated with natural ponds and is particularly vulnerable to landscape fragmentation due to urbanisation (Emaresi et al., 2011; Van Buskirk, 2012), were only recorded in the Bakony Mountains, a protected forest in western Hungary. *Salamandra salamandra* can persist in low-density urban areas provided there is sufficient forest nearby. Accordingly, it was reported from a garden pond adjacent to a large forested area (Bükk Mountains in northern Hungary) and interestingly, a population exists even at the fringe of the capital city Budapest (Kiss et al., 2022a).

Conversely, occurrences of odonates and birds were positively associated with urbanisation in general. These results might seem counterintuitive at first, although there is evidence that in certain cases urban areas are able to maintain higher odonate species diversity than agricultural lands, due to a network of public, stormwater, or garden ponds that can foster the migration of species (Goertzen & Suhling, 2019; Willigalla & Fartmann, 2012). However, other studies showed that species richness of odonates increased from the city center to more rural areas, probably due to increasing habitat availability (more water bodies in the rural areas than in cities) (Holtmann et al., 2018; Willigalla & Fartmann, 2012). Here, we only recorded the presence of odonates as a group. Therefore, the observed positive association could also be explained by a few highly tolerant species occurring in urbanised landscapes, but more detailed follow-up studies will be necessary to link this pattern to species identities and biodiversity. The same might be true for birds, resulting from the presence of more generalist bird species such as blackbirds or sparrows, which are common even in heavily urbanised landscapes (Callaghan et al., 2019; Mennechez & Clergeau, 2006). Furthermore, birds are more likely to visit garden ponds, puddles, or fountains, in urban areas, where other natural water sources are scarce (Tryjanowski et al., 2022), which could have contributed to the pattern we found.

### 4.2. The most influential pond features

Pond age was overall the most important pond feature shaping the occurrence of animal taxa. Newly installed ponds (0-1 years) were less occupied by amphibians and their tadpoles, odonates and birds, which can be due to the lack of vegetation and sediment that could offer hiding and breeding places to aquatic and semi-aquatic animal taxa (Biggs et al., 2005; Huikkonen et al., 2020; Soukup et al., 2022). Additionally, colonisation time might be another factor as animals need time to find and occupy newly installed ponds in the urban landscape (Cayuela et al., 2020; Coccia et al., 2016; Unglaub et al., 2021). For example, many odonate species have longer life cycles, so adult life stages may only reach higher densities at older ponds (Hassall & Thompson, 2008; Stoks & Córdoba-Aguilar, 2012), hence pond age may increase encounter probabilities with them. Most amphibians and their tadpoles were significantly associated with older ponds, corresponding to a longer period for species to locate and colonise these ponds (Moor et al., 2022). Among them, *Rana arvalis* was an exception, being strongly associated with newly installed ponds that this species can potentially colonise while moving relatively long distances through the landscape (Brzeziński & Mętrak, 2014). Many amphibian species prefer well-vegetated or large ponds (Arntzen et al., 1997; Péntek et al., 2017; Rannap et al., 2013), which was also supported by our results as *Bombina bombina, Pelobates fuscus, Triturus dobrogicus* were positively associated with aquatic vegetation. Aquatic vegetation was also positively associated with tadpoles, odonates, and bird presence which indicates the habitat structuring rola of aquatic vegetation that benefits biodiversity (Ansari et al., 2017; Law et al., 2019). Unexpectedly, rubber pond liner had a significantly positive effect on the presence of amphibians and their tadpoles, and also on odonates, birds, and wading birds. This may have resulted from the generally larger size of the rubber-lined ponds as shown by an earlier study (Hamer et al., 2024).

### 4.3. Management of garden ponds

The use of chemicals, such as pesticides, fertilizers, and algaecides is common around ponds in agricultural and recreational settings like golf courses, but they can also be applied in private gardens to improve soil and water quality or control biological processes (Boyd & Massaut, 1999; Oertli & Parris, 2019). Even though pesticides are intended to target specific pests, they can affect other non-target organisms via overland runoff, potentially impacting nearby habitats (Rumschlag et al., 2020; Yamamuro et al., 2019). However, private gardens, including those with ponds, are characterized by relatively lower levels of pesticide use, due to which they can even serve as refuges for pollinators (Hall et al., 2017; Siviter et al., 2023). Furthermore, unlike public ponds that often receive polluted surface runoff (Ivanovsky et al., 2018; Osman et al., 2019), garden ponds are usually isolated from such events. As a result, fertilizers and insecticides applied elsewhere in the garden have limited potential to contaminate the ponds. Additionally, insecticides are also typically applied on garden crops, and therefore they represent minimal risk to garden pond fauna, as these are usually not applied on aquatic plants such as water lilies and sedges.

On the other hand, the direct application of algaecides was a common practice among Hungarian garden pond owners according to our survey (reported from 33% of the ponds). Our findings imply a negative effect of algaecides on amphibians and their tadpoles. At the same time, our multivariate analysis showed a strong association between algaecide use and the presence of fish, and both were negatively aligned with almost all amphibian species. Hence the negative effect of algaecides on the presence and reproduction of amphibians might be also linked to the presence of fish in the same ponds. High nutrient inputs in the form of fish food and a low abundance of aquatic macrophytes, which are competitors of phytoplankton (Zeng et al., 2022), contributes to high algal production that could subsequently lead to algaecide application in these ponds. While we did not collect specific information on the exact types of algaecide used, several respondents, on their own accord, indicated a specific brand of algae managing substance which uses a natural approach (i.e. microbial decomposition) rather than artificial chemicals, which points more at the indirect link through fish.

### 4.4. Invasion risk of the introduced animals and the role of fish

Based on direct communication with the pond owners, exotic animals are frequently introduced to garden ponds for ornamental purposes (e.g. goldfish, koi) (Hassall, 2014). According to our results, fish in general were the most frequently introduced animals, being consequently present in 84% of ponds. In comparison to our results, studies conducted in the UK and the Netherlands reported that only ∼35-32% of the garden pond owners stocked their ponds with fish (Loram et al., 2011; Peeters et al., 2023). Besides fish, turtles and several invertebrates (snails, mussels, and crayfish) were also reported. Previous studies found that aquaria and garden ponds may act as sources of invasions (C. Chucholl, 2013; Patoka et al., 2014; Soes et al., 2011; Van Den Neucker et al., 2017), for example in the case of the red-eared terrapin (*Trachemys scripta elegans*) (Tietz et al., 2023), or signal crayfish (*Pacifastacus leniusculus*) (F. Chucholl et al., 2021). In this sense, without the indispensable knowledge and education about invasive species, garden pond owners and garden centres can pose a threat to native biodiversity. This highlights the importance of outreach activities targeting pond owners, encouraging more conscious management choices.

We also found that the introduction of fish was negatively associated with all recorded amphibian species. Fish are important predators of all amphibian life stages and a negative relationship between fish and amphibian populations has been demonstrated for many European amphibians (Hartel et al., 2007; Van Buskirk, 2012). But despite the introduction of fish into 84% of ponds, amphibian reproduction was observed in 62% of the ponds, indicating that some species are able to coexist with fish (Hamer et al., 2021; Kloskowski et al., 2020). Aquatic vegetation can provide refugia for amphibians from fish predation (Hartel et al., 2007), which could be also observed in our dataset, where well-vegetated ponds supported a higher co-existence of fish and amphibians (73.5%) than poorly vegetated ponds (54%). As the introduction of fish was a significant predictor of garden pond design, within-pond features, and management practices in these ponds (Hamer et al., 2024), this result implies a potential conflict between creating ponds with the main aim of keeping ornamental fish and their potential to support native biodiversity. However, our results imply that planting dense aquatic vegetation can mitigate the negative effect of fish on amphibians in garden ponds.

Also, wading birds were frequently (24.3%) reported from the ponds, the majority of which were stocked with fish (18.1% of the ponds in our survey). Sightings in this group included grey herons, which regularly utilise garden ponds for foraging on fish (Jakubas & Manikowska, 2011). This group represents a source of human-nature conflict as, according to anecdotal evidence and comments given as part of the responses in our survey, they can damage ponds by puncturing pond liners and prey upon ornamental fish.

### 4.5. Caveats and relevance of using citizen science data in garden pond surveys

We identified a number of caveats related to the use of citizen science data as part of our study. Due to reliability issues, we lost 9.7% of the survey responses in the first step of the validation (excluding duplicates and incomplete data). In the case of amphibian species and bird types, even with our best efforts to validate the dataset, there is a chance of misidentification. To overcome this, we collected data on taxa that can easily be identified, and are considered typical groups representing pond biodiversity (Oertli et al., 2002). We also provided an identification key for amphibian species (MME 2022. https://www.mme.hu/en/node/4440) along with our survey. Another unexpected caveat was the unit of pond size for example, where some respondents did the conversion of centimeter to meter or meter to centimeter incorrectly. In these cases, we manually checked the pond size with Google Maps based on GPS coordinates. Although we can not compare our questionnaire data to earlier or other regional datasets as a reference, the high number of ponds in our study provides robust data for the general patterns observed in these poorly studied secondary habitats. These could later serve as important baselines for more environmentally friendly management activities, being more beneficial for the local pond biota.

Increasing pressure for urban development poses a significant threat to blue-green infrastructure, reducing its extent and complicating its management (Ancillotto et al., 2019). Citizen science can provide important means for efficient large-scale biodiversity monitoring, particularly in cases where access to private properties is limited, as was the case in our study. The great potential of the approach is well-illustrated by the high level of engagement in the present survey, with over 800 completed forms. Also, citizen science emerges as a powerful tool for urban planning, as it can contribute to gathering valuable data on urban biodiversity and utilise it for more efficient conservation strategies. For instance, it could help urban planning directly by identifying hotspots of aquatic biodiversity or critical areas for the conservation of key groups like amphibians in urban environments. Amphibians are one of the central targets for pond creation projects in Europe (De Necker et al., 2024) and there is increasing evidence that urban ponds, including garden ponds, can contribute to their conservation in the urban landscape by offering valuable secondary habitats and potentially acting as stepping stones between larger habitats (Butterworth et al., 2025; Hamer et al., 2021; Hamer & Parris, 2011; Kiss et al., 2022b). Citizen science data could also provide a feasible way to assess the green infrastructure of cities and could be used for the long-term monitoring of the potential effects of management activities on urban biodiversity (Callaghan et al., 2020). In our study, we collected a large-scale biodiversity dataset of garden ponds, which could potentially help to identify ponds or larger areas hosting protected species. Such programs may help authorities to make informed decisions on planning and managing the blue-green infrastructure to support the biodiversity of protected species, for example by creating new ponds in parks near to garden ponds with the known occurrence of target species. Information on the location of these new connecting elements of aquatic habitat networks can aid the formation of corridors increasing the connectivity between larger aquatic habitats in cities (Hyseni et al., 2021). Without the active engagement of citizens, such information would not be available for city planners. Finally, active citizen engagement not only contributes to large-scale data collection but it is also a tool for environmental education. Participants may become more conscious of environmental issues and their role in it which might lead to more active engagement in supporting blue-green infrastructure development.

## 5. Conclusions

To date, there has been limited information available on the biodiversity of garden ponds worldwide. In this study, we present the first large-scale dataset evaluating the effects of urbanisation, local features, and management practices on garden pond biodiversity using responses from 753 pond owners in Hungary. Our results highlighted the importance of garden ponds for urban biodiversity and conservation, as they serve as secondary habitats for multiple animal taxa in urban environments. This implication can be directly communicated to the public while promoting pond installations in urban areas, where the importance of aquatic vegetation can be emphasised. These activities, however, should also highlight relevant facts about invasive species and the most suitable local features and management types to maximise local biodiversity. For effective biodiversity conservation in urban areas, where scarce natural habitats are under constant pressure from degradation and mismanagement, public engagement is becoming increasingly valuable. Citizen science not only provides essential tools for gathering data but could be utilised to further engage citizens in their crucial role as pond managers.

## Web references

https://mypond.hu/en/ - Last accessed: 2023.11.23.

https://herpterkep.mme.hu/index.php?lang=en - Last accessed: 2023.11.23.

## List of appendices

**Table S1.**
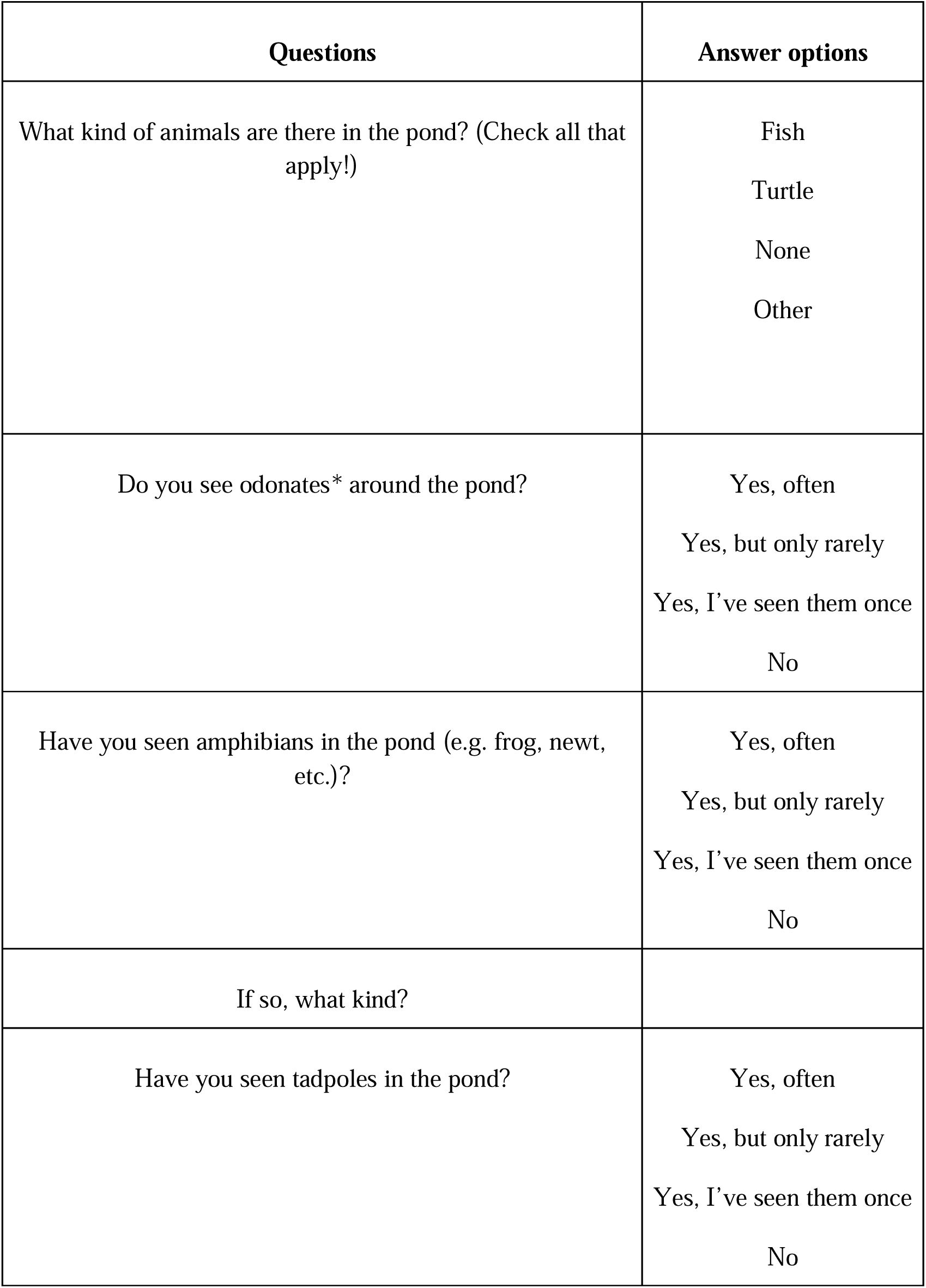
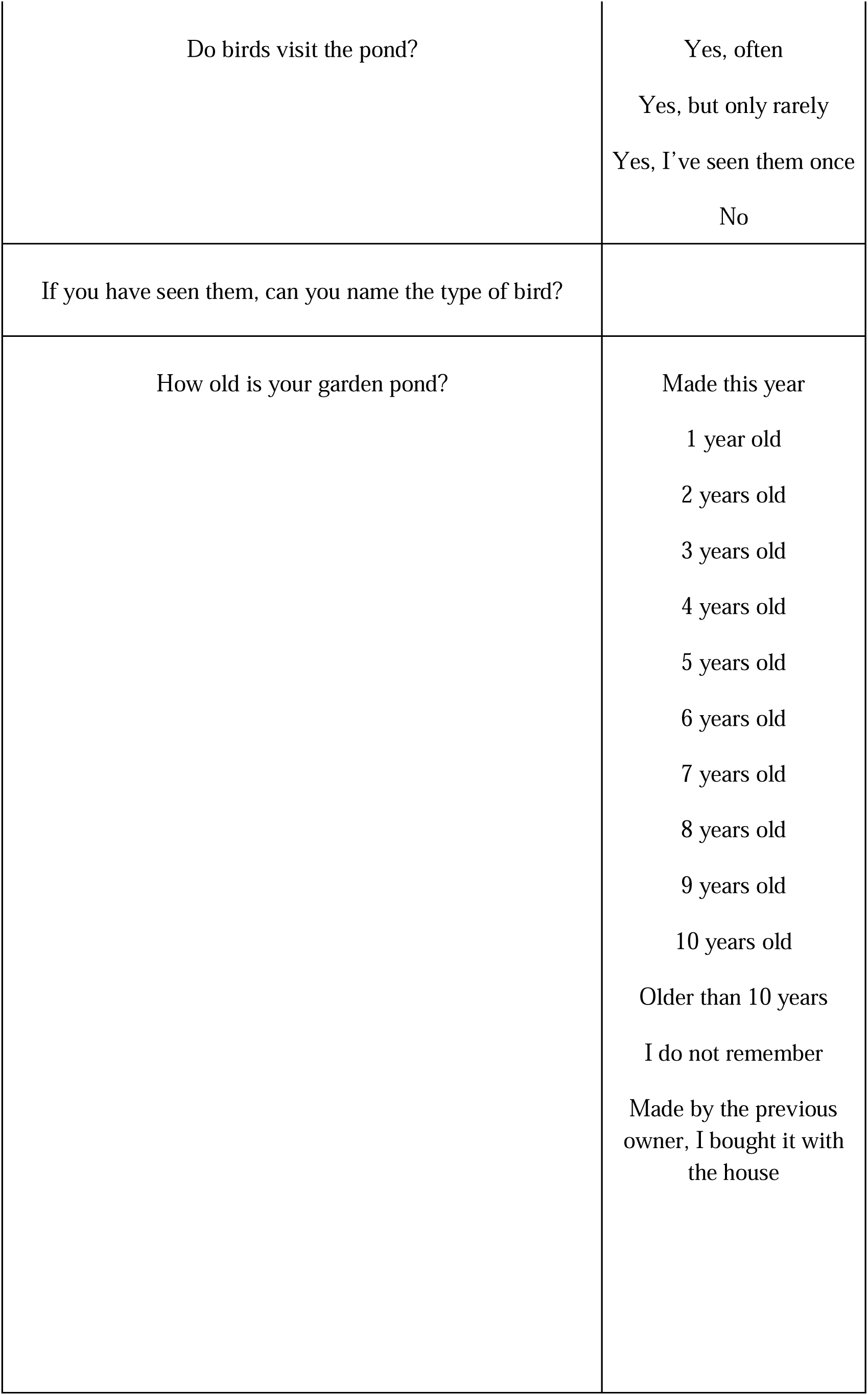
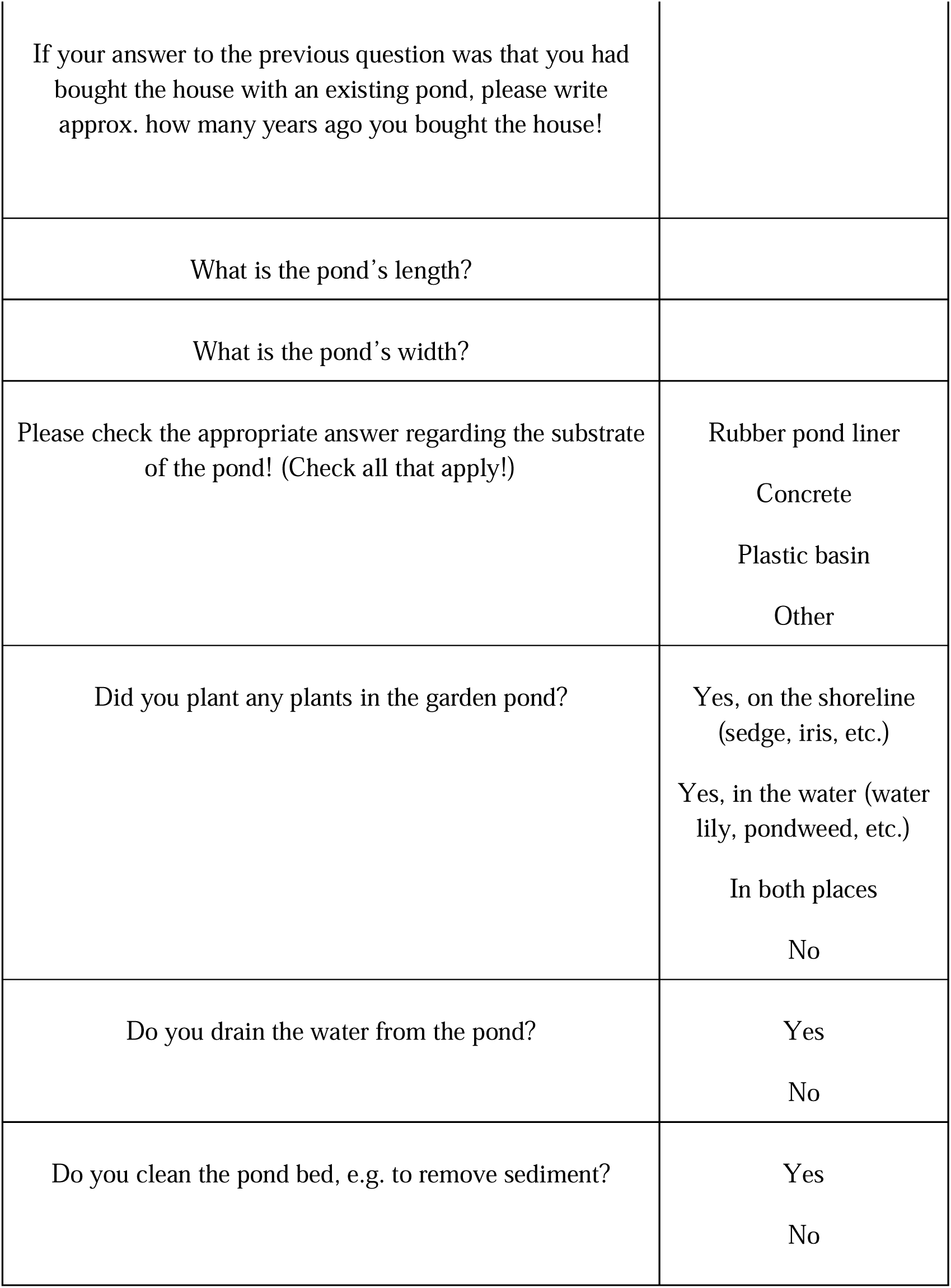
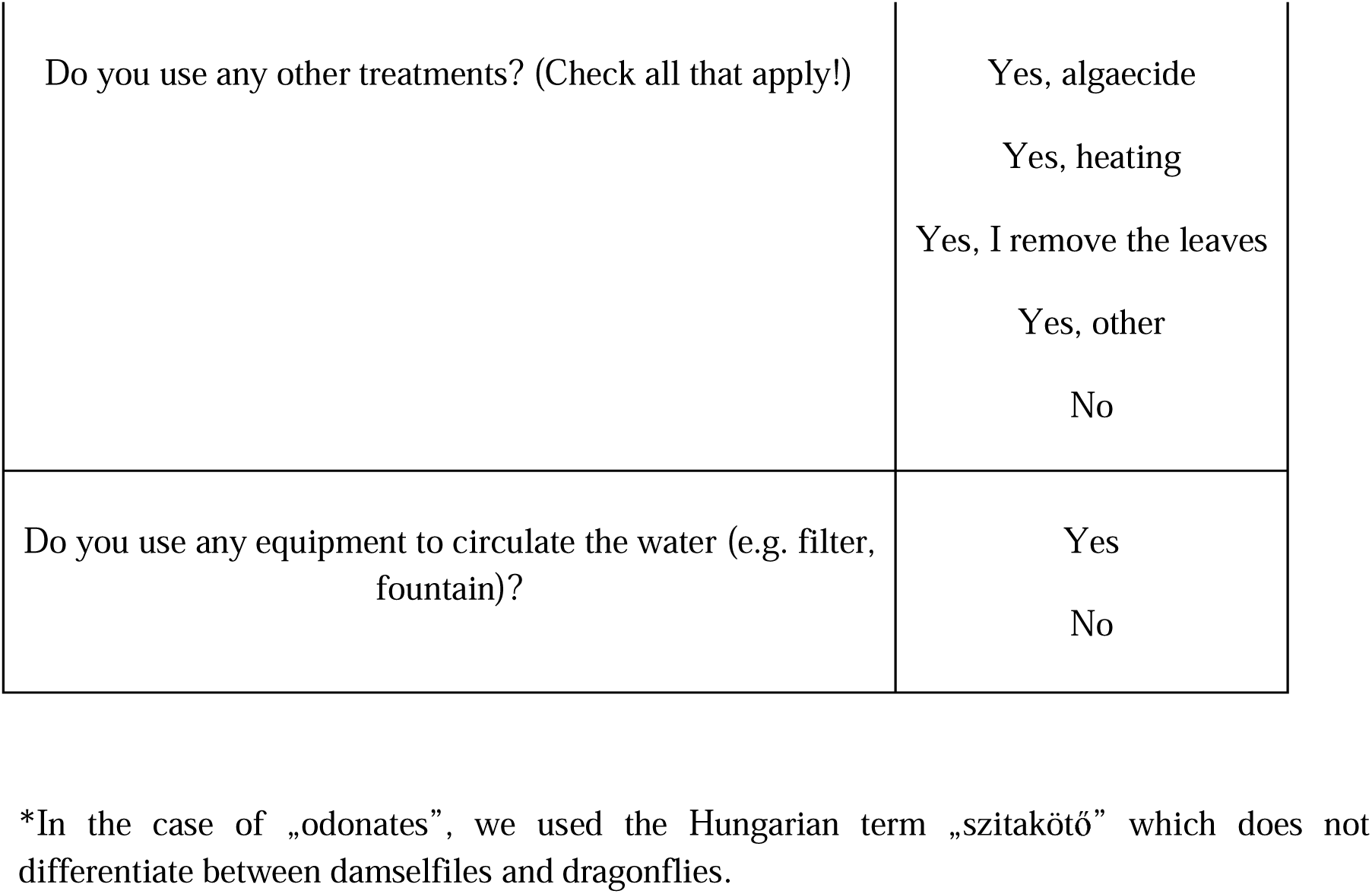
Questions of the “MyPond” online survey.

| Questions | Answer options |
| --- | --- |
| What kind of animals are there in the pond? (Check all that apply!) | Fish<br>Turtle<br>None<br>Other |
| Do you see odonates* around the pond? | Yes, often<br>Yes, but only rarely<br>Yes, I’ve seen them once<br>No |
| Have you seen amphibians in the pond (e.g. frog, newt, etc.)? | Yes, often<br>Yes, but only rarely<br>Yes, I’ve seen them once<br>No |
| If so, what kind? |  |
| Have you seen tadpoles in the pond? | Yes, often<br>Yes, but only rarely<br>Yes, I’ve seen them once<br>No |
| Do birds visit the pond? | Yes, often<br>Yes, but only rarely<br>Yes, I've seen them once<br>No |
| If you have seen them, can you name the type of bird? |  |
| How old is your garden pond? | Made this year<br>1 year old<br>2 years old<br>3 years old<br>4 years old<br>5 years old<br>6 years old<br>7 years old<br>8 years old<br>9 years old<br>10 years old<br>Older than 10 years<br>I do not remember<br>Made by the previous owner, I bought it with the house |
| If your answer to the previous question was that you had bought the house with an existing pond, please write approx. how many years ago you bought the house! |  |
| What is the pond's length? |  |
| What is the pond's width? |  |
| Please check the appropriate answer regarding the substrate of the pond! (Check all that apply!) | Rubber pond liner<br>Concrete<br>Plastic basin<br>Other |
| Did you plant any plants in the garden pond? | Yes, on the shoreline (sedge, iris, etc.)<br>Yes, in the water (water lily, pondweed, etc.)<br>In both places<br>No |
| Do you drain the water from the pond? | Yes<br>No |
| Do you clean the pond bed, e.g. to remove sediment? | Yes<br>No |
| Do you use any other treatments? (Check all that apply!) | Yes, algacide<br>Yes, heating<br>Yes, I remove the leaves<br>Yes, other<br>No |
| Do you use any equipment to circulate the water (e.g. filter, fountain)? | Yes<br>No |

\*In the case of „odonates”, we used the Hungarian term „szitakötő” which does not differentiate between damselflies and dragonflies.

**Table S2.** Summary statistics of the Moran’s I spatial autocorrelation tests used for each GLM model listed in Table S3.

| Group | Observed | Expected | sd | p value |
| --- | --- | --- | --- | --- |
| Amphibians | 0.007 | -0.001 | 0.009 | 0.355 |
| Tadpoles | 0.002 | -0.001 | 0.009 | 0.461 |
| Odonates | 0.011 | -0.001 | 0.009 | 0.196 |
| Birds | 0.011 | -0.001 | 0.009 | 0.196 |
| Ducks | 0.001 | -0.001 | 0.009 | 0.765 |
| Other birds | 0.005 | -0.001 | 0.009 | 0.464 |
| Wading birds | 0.012 | -0.001 | 0.009 | 0.167 |

**Table S3.** Summary statistics of the best-fit logistic regression models (based on AIC) testing the relationship between the prevalence of animal groups and pond features and management.

|  |  |  |  |  |  |  | **Odds ratio results** |  |
| --- | --- | --- | --- | --- | --- | --- | --- | --- |
| **Group** | **Final model elements** | **Estimate** | **Std.Error** | **z value** | **p value** | **F value** | **Odds ratio** | **Confidence Interval (95%)** |
| **Amphibians** | Age 0-1 | -1.261 | 0.258 | -5.262 | <0.001 | 28.830 | 0.256 | 0.154-0.426 |
|  | Area | 0.036 | 0.011 | 3.310 | 0.001 | 19.426 | 1.036 | 1.016-1.061 |
|  | Rubber pond liner | 0.765 | 0.266 | 2.871 | 0.004 | 8.644 | 2.150 | 1.269-3.618 |
|  | Natural substrate | 14.442 | 528.959 | 0.027 | 0.978 | 4.195 | 1.872E+06 | 0.000-2.157E+75 |
|  | Algaecide | -0.421 | 0.208 | -2.024 | 0.043 | 4.368 | 0.656 | 0.436-0.988 |
|  | Urbanisation | -1.956 | 0.436 | 4.487 | <0.001 | 23.693 | 0.141 | 0.058-0.326 |
|  | Wetland coverage | 3.535 | 2.207 | 1.601 | 0.109 | 3.160 | 34.300 | 0.621-3810.007 |
| **Tadpoles** | Age 0-1 | -1.045 | 0.233 | 4.483 | <0.001 | 19.557 | 0.351 | 0.221-0.553 |
|  | Area | 0.014 | 0.004 | 3.312 | 0.001 | 14.316 | 1.014 | 1.006-1.024 |
|  | Concrete substrate | 0.471 | 0.321 | 1.468 | 0.142 | 2.101 | 1.601 | 0.862-3.048 |
|  | Rubber pond liner | 0.692 | 0.234 | 2.954 | 0.003 | 8.344 | 1.997 | 1.264-3.171 |
|  | Algaecide | -0.631 | 0.167 | 3.756 | <0.001 | 13.412 | 0.532 | 0.382-0.739 |
|  | Shoreline vegetation | 1.120 | 0.481 | 2.326 | 0.020 | 5.723 | 3.066 | 1.243-8.407 |
|  | Aquatic vegetation | 0.457 | 0.222 | 2.056 | 0.040 | 3.987 | 1.580 | 1.020-2.446 |
|  | Urbanisation | -1.236 | 0.315 | -3.916 | <0.001 | 15.010 | 0.290 | 0.155-0.536 |
| **Odonates** | Age 0-1 | -1.866 | 0.438 | -4.257 | <0.001 | 17.374 | 0.154 | 0.065-0.368 |
|  | Rubber pond liner | 0.920 | 0.465 | 1.979 | 0.048 | 3.785 | 2.510 | 0.969-6.118 |
|  | Shoreline vegetation | 16.047 | 1102.637 | 0.015 | 0.988 | 4.847 | 9.321E+06 | 3.921E-19-9.744E+94 |
|  | Aquatic vegetation | 1.281 | 0.461 | 2.779 | 0.005 | 7.443 | 3.599 | 1.423-8.833 |
|  | Urbanisation | 1.236 | 0.748 | 1.651 | 0.099 | 2.891 | 3.443 | 0.799-15.355 |
| **Birds** | Age 0-1 | -1.866 | 0.438 | -4.257 | <0.001 | 17.374 | 0.155 | 0.065-0.368 |
|  | Rubber pond liner | 0.920 | 0.465 | 1.979 | 0.048 | 3.785 | 2.510 | 0.9697-6.118 |
|  | Shoreline vegetation | 16.047 | 1102.637 | 0.015 | 0.988 | 4.847 | 9.321E+06 | 3.921E-19-9.744E+94 |
|  | Aquatic vegetation | 1.281 | 0.461 | 2.779 | 0.005 | 7.443 | 3.599 | 1.423-8.833 |
|  | Urbanisation | 1.236 | 0.748 | 1.651 | 0.099 | 2.891 | 3.443 | 0.799-15.355 |
| **Ducks** | Urbanisation | -0.667 | 0.447 | -1.492 | 0.136 | 2.209 | 0.512 | 0.212-1.235 |
| **Other birds** | Cleaning | 0.708 | 0.328 | 2.158 | 0.031 | 4.754 | 2.031 | 1.073-3.925 |
|  | Urbanisation | 1.239 | 0.612 | 2.023 | 0.043 | 4.131 | 3.454 | 1.044-11.638 |
| **Wading birds** | Area | -0.006 | 0.004 | -1.513 | 0.131 | 2.673 | 0.993 | 0.984-1.001 |
|  | Rubber pond liner | 1.011 | 0.424 | 2.382 | 0.017 | 6.579 | 2.751 | 1.254-6.716 |
|  | Plastic<br>substrate | 1.104 | 0.445 | 2.479 | 0.013 | 6.539 | 3.01 | 1.288-7.513 |
|  | Draining | -0.284 | 0.198 | -1.433 | 0.152 | 2.087 | 0.752 | 0.506-1.105 |
|  | Circulation | -0.341 | 0.209 | -1.632 | 0.102 | 2.605 | 0.711 | 0.473-1.076 |

